# Whole-proteome Tree of Insects: Grouping and phylogeny without sequence alignment

**DOI:** 10.1101/2020.07.11.198689

**Authors:** JaeJin Choi, Byung-Ju Kim, Sung-Hou Kim

**Author notes:** Corresponding author: Sung-Hou Kim, 419 Latimer Hall, University of California, Berkeley CA, 94720, USA. e-mail addresses of co-authors: JJC; BJK.

## Abstract

An “organism tree” of insects, the largest and most species-diverse group of all living animals, can be considered as a conceptual tree to capture a simplified narrative of the complex evolutionary courses of the extant insects. Currently, the most common approach has been to construct a “protein tree”, as a surrogate for the organism tree, by Multiple Sequence Alignment (MSA) of highly homologous regions of a set of select proteins to represent each organism. However, such selected regions account for a very small fraction of the whole-proteome of each organism.

Information Theory provides a method of comparing two sets of *all* proteins, two whole-proteomes, without MSA: By treating each whole-proteome sequence as a “book” of amino acid alphabets, the information contents of two whole-proteomes can be quantitatively compared using the text comparison method of the theory, without sequence alignment, providing an opportunity to construct a “whole-proteome tree” of insects as a surrogate for an organism tree of insects.

A whole-proteome tree of the insects in this study shows that: (a) all the founders of the major groups of the insects have emerged in an explosive “burst” near the root of the tree, (b) the most basal group of all the insects is a subgroup of Hemiptera consisting of aphids and psyllids, and (c) there are other notable differences in the phylogeny of the groups compared to those of the recent protein trees of insects.

## Introduction

### Sequence-alignment-based “protein trees”

An “organism tree” of insects can be considered as a practically useful narrative to convey a simplified evolutionary relationship among the insects. However, it is a conceptual tree that cannot be experimentally validated. Thus, it is expected that the effort will continue to find one or more “surrogate trees**”** derived from various descriptors of the characteristics associated with each insect and to find improved methods to estimate evolutionary distances from the divergence of the descriptors under as few subjective assumptions as possible at the time of investigation.

At present, the best descriptor of an insect as an organism is its entire whole-genome sequence information, from which whole-proteome sequence can be derived. However, for several decades, due to the technical difficulties and high cost of whole genome sequencing, and to the difficult task of comparing unaligned sequences, the most practically feasible and common approach to construct a surrogate tree has been to construct a Multiple Sequence Alignment (MSA)-based “protein tree” under a few important, but debatable assumptions: (a) a set of regions with high homology selected among each of homologous proteins may have enough information to represent a whole organism, and (b) the divergence of certain characteristics, most commonly, point substitution rates within each MSA aligned-regions may be a reasonable measure to represent the evolutionary divergence/distances among the whole organisms, without considering possible evolutionary roles of all other proteins without high homologous regions among them and of other mutational events including absence/presence of proteins.

Such “alignment-based” protein trees represent the evolutionary phylogeny of the selected regions of the selected proteins, but not full characteristics of all proteins, let alone the whole organisms, because the aligned regions account, in general, for a very small fraction of all proteins (Pace 2009).

### Information-theory-based (“alignment-free”) “whole-proteome trees”

This situation has since changed significantly in two important aspects: (a) During last decades, a large number (over 134 species, mostly insects, as of 2020) of whole-genome sequences of extant insect species have been accumulating in public databases, and (b) Information Theory, developed to analyze linear electronic signals, was found to be adaptable to analyze other linear information, such as natural languages and genomic information, without sequence alignment (“alignment-free”) (Zielezinski et al. 2017; Blaisdell 1986). In this approach, the whole content, not selected portions of the whole content, of a whole-proteome sequence, can be described by “*n*-Gram” or “*k*-mers” (Zielezinski et al. 2017). *N*-Gram of a whole-proteome is the collection of all overlapping short subsequences of length *n*, and it contains all information necessary to reconstruct the original sequence. Furthermore, the information divergence (difference) between two *n*-Grams can be estimated by, for example, Jensen-Shannon divergence (JSD) without alignment of the whole proteome sequences (Lin 1991). Such approach has been widely tested and validated for comparing texts and books of natural languages for latent semantic analysis since 1990s (Deerwester et al. 1990) and gene sequences consisting of coding and non-coding regions as well as amino acid sequences since 1986 (Blaisdell 1986).

Some of these validated methods have been adapted and optimized to handle whole-proteome sequences in Feature Frequency Profile (FFP) method (Sims et al. 2009). Since there is no “golden standard” for a phylogenetic evolutionary tree of a group of organisms that can be experimentally validated, the FFP method has been tested using 26 books in English alphabets from diverse authors and genres, after removing all spaces and delimiters as well as author names, book titles, headers, footers, etc. In general, the method performed well in grouping the “books” by the genre and authors (see Fig. 1 of Sims et al. 2009). In a recent bench-marking studies of 24 Alignment-free methods, FFP method was ranked among the top 5 best-performing tools for phylogeny prediction based on the input data of assembled whole genome sequences (Zielezinski et al. 2019).

**Fig 1:**
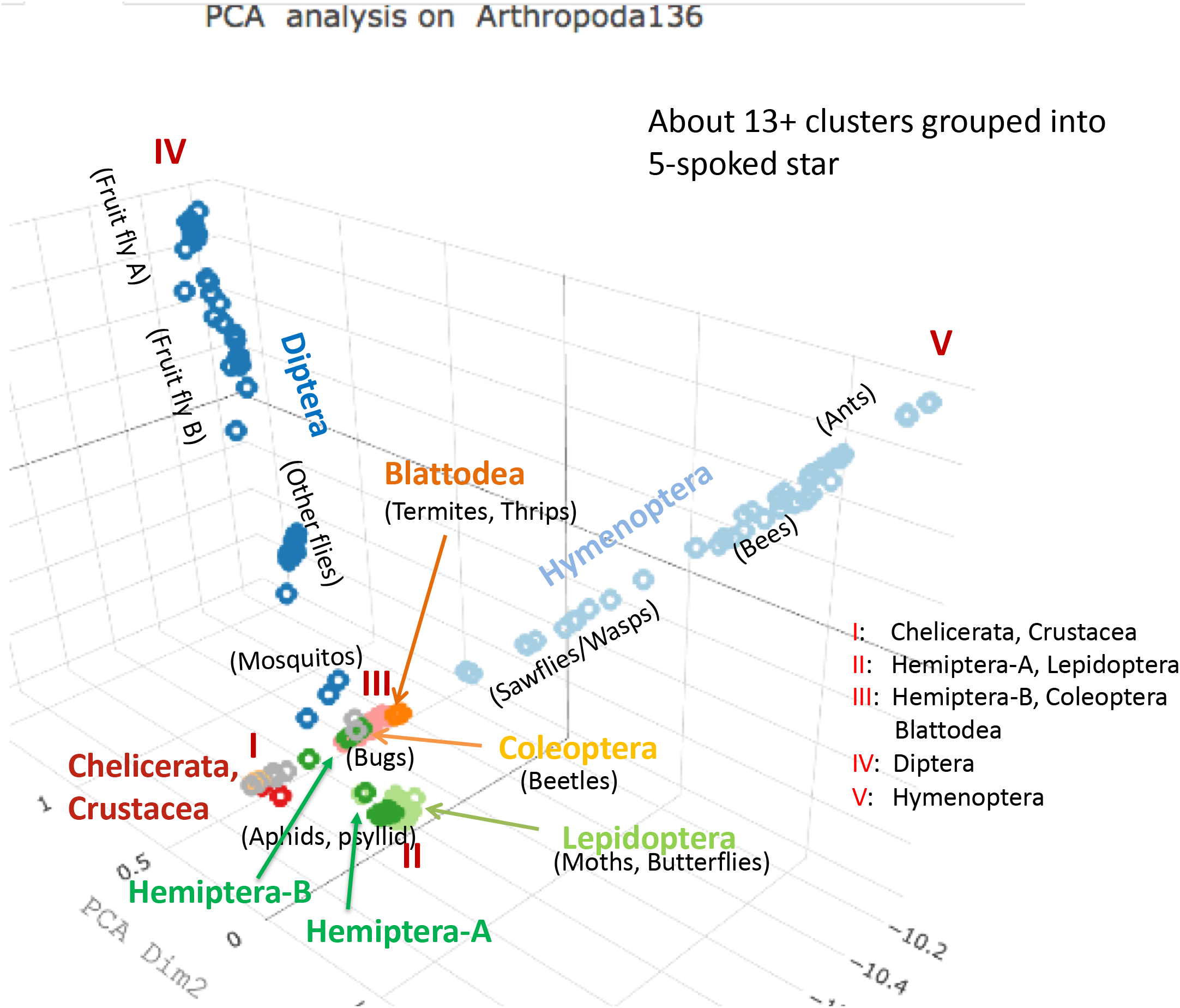
Unsupervised clustering (grouping) of 134 extant arthropods (123 insects plus 11 non-insect arthropods) by classical PCA. Classical PCA plotted for the three principal axes reveals about 5 large clusters arranged in 5-spokes. Two long spokes (IV and V) corresponds to all the members of Diptera and Hymenoptera, respectively. The remaining three short spokes (I, II, and III) correspond to: Members of Chelicerata and Crustacea in spoke I; those of Hemipters-A and Lepidoptera in spoke II, and those of Hemiptera-B, Coleoptera, and Blattodea in spoke III.

In the FFP method, whole-genome/whole-proteome information is used under the assumptions very different from the protein trees: (a) whole-proteome sequence of an organism represents the organism better than the collection of short regions of highly homologous sequence from a set of selected genes/proteins used in the protein trees and (b) a combination of *all types of mutations*, such as point substitution, insertion/deletion of various length, recombination, duplication, transfer or swapping of genes etc., contribute to the evolutionary processes of the organisms, rather than only point substitution rates in the protein trees. Thus, whole-proteome tree may provide an independent view of the evolutionary relationship among the insect organisms.

### Experiences from earlier whole-proteome trees

In the last decade, we have tested and optimized the protocols for building whole-proteome trees using various different populations such as Bacteria and Archaea Domains (Jun et al. 2010), Fungi Kingdom (Choi & Kim 2017) and, most recently, all three Domains at a deep phylogenic level (Choi & Kim 2020). From these studies we have learned that: (a) among three types of the trees (whole-genome DNA tree, whole-transcriptome RNA tree, and whole-proteome amino-acid tree) the whole-proteome trees produce the most topologically stable trees; (b) for a give group of organisms, the optimal length of the sequence strings to be used in FFP method can be empirically determined; and (c) cumulative genomic divergence (CGD) is a useful and computable quantity for the point of the emergence for the founders of a group in the “evolutionary progression scale”.

In this study, we optimized various parameters and protocols specifically for the population of insects and present a view of the whole-proteome tree based on whole-proteome sequences of 134 diverse arthropod species (123 insects plus 11 non-insect arthropods), available in the NCBI database (O’Leary et al. 2016), and discuss its implications to phylogenic aspects of insect evolution.

## Results

To compare the current protein trees with our whole-proteome Tree of Insects (ToIn) we chose two recent and very comprehensively-analyzed protein trees: The first one is the recent “alignment-based” tree of 144 insect taxa based on 1,478 single-copy protein-coding nuclear genes (Fig. 1 of Misof et al. 2014). The second is the tree for 76 arthropod taxa (Fig. 2 of Thomas et al. 2020) based on up to 4,097 single copy protein-coding genes. In both cases, the number of aligned genes used are a small fraction of about 10,000 to 31,000 genes among their study insects. Both protein trees agree with each other, in general, on the branching order of the Order groups of the respective populations, and on similar time spread of the emergence of the founders of the groups in chronological time scale estimated based on available fossils and calibration methods under various assumptions.

**Fig 2:**
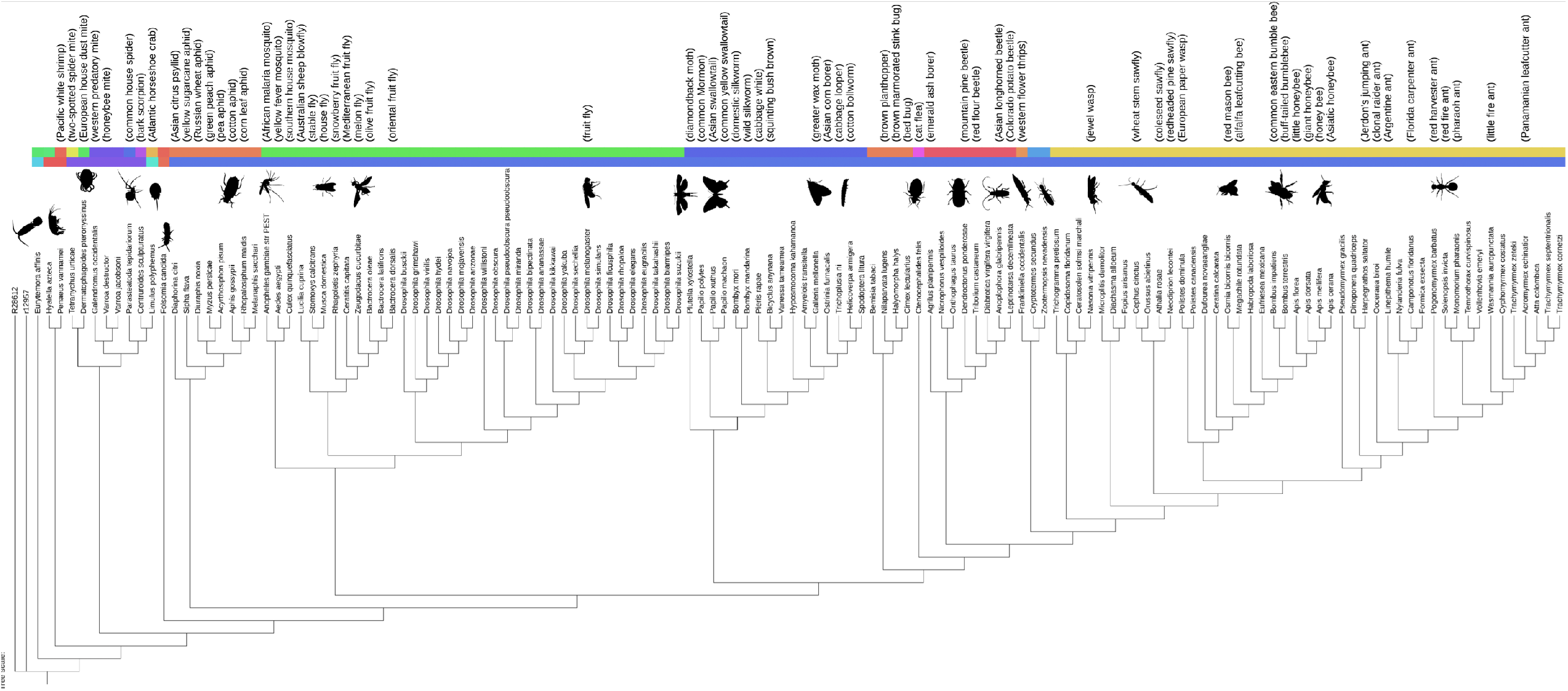
Topology of the linear representation of whole-proteome Tree of Insects (ToIn) The colors of the first (inner) colored-band distinguish organisms in different Classes, and those of the second (outer) band among different Orders (the names of different color-bands are shown in Fig. 3A). Scientific names and common names, when available, of each organism are also listed. The silhouettes of sampled organisms are shown next to their names. To emphasize the clading pattern, all branch-lengths are ignored. The first two items refer to two members of the outgroup (see Outgroup in Discussions and Implications) constructed by shuffling (Knuth 1973; Fisher & Yates 1948) the whole-proteome sequences of the two arthropods. The visualization of the ToIn was made using iTOL (Letunic & Bork 2019).

In comparing our whole-proteome ToIn to these two protein trees, we focus on two aspects separately: grouping patterns and phylogeny of the groups. For the former, we use two methods: First, we cluster our study population by several unsupervised clustering algorithms using only the distances estimated from the “divergence” among whole-proteome FFPs with no explicit constraints of the presence of the common ancestor(s) or specific evolutionary models (see Construction of whole-proteome Tree of Insects in Materials and Methods). We then ask whether the “clustering pattern” is similar to the “clading pattern” in the protein trees and in our whole-proteome ToIn, recognizing that the both tree constructions assume the constrains of the common ancestor(s) and specific evolutionary models. For the phylogeny of the groups, we compare the order of branching of the groups and their emergence points on the evolutionary progression scale in our tree and in chronological scale in the protein trees (see “Cumulative Genomic Divergence (CGD)” as “Evolutionary Progression Scale” in Materials and Methods).

### A. Demographic grouping pattern by clustering and clading

#### Clustering

We have tested the grouping pattern of the insects by several unsupervised clustering algorithms, such as Principal Component Analysis (PCA), Multi-Dimensional Scaling, and t-Distributed Stochastic Neighbor Embedding (t-SNE) (R Core Team 2016; v.d. Maaten & Hinton 2008), all of which can be accomplished from the same starting “distance” matrix constructed using the divergence of whole-proteome sequences, as calculated by JSD (Lin 1991) of FFPs, among all pairs of the study organisms. All three clustering methods showed the clustering pattern compatible with the current grouping of arthropods with common and scientific names, mostly based on morphological characteristics. Figure 1, a classical PCA clustering, which is very similar to that of MDS method, shows that all our study population are distributed into 5 “spokes”. Two long spokes (IV and V) corresponds to all the members of Diptera and Hymenoptera of Insecta Class, respectively. The remaining three short spokes (I, II, and III) correspond to: Members of Chelicerata and Crustacea of non-insect arthropods in spoke I; those of Hemipters-A and Lepidoptera of Insecta Class in spoke II, and those of Hemiptera-B, Coleoptera, and Blattodea of Insecta Class in spoke III. The most noticeable difference with the grouping in the current protein trees (Misof et al. 2014; Thomas et al. 2020) is that Hemiptera is split into two separate groups, labeled in this study as Hemiptera-A and -B. Another clustering by t-SNE (see Supplemental information, Fig. S1) also shows a similar split of Hemiptera into two, which is unexpected, because the assumptions and algorithms in t-SNE and PCA are completely different.

#### Clading

As a second method of the grouping, we use the clading pattern of the organisms in our whole-proteome ToIn. Figure 2 shows the topology of the ToIn, constructed using Neighbor-Joining method implemented in BIONJ (Saitou & Nei 1987; Gascuel 1997). In this study, we use the divergence of whole-proteome sequences of two organisms as the estimates for the evolutionary distances between them, as calculated by JSD of pair-wise FFPs at an optimal Feature length (see Construction of whole-proteome Tree of Insects in Materials and Methods and Supplementary information Fig. S2). We also assume an evolutionary model of Maximum Parsimony (minimum evolution) in a way that the chosen neighbors to be joined are those that minimize the total sum of branch-lengths at each stage of step-wise joining of neighbors starting from a star-like tree (Saitou & Nei 1987). The tree shows that most of the clusters in Fig.1 can be identified among the clades in the ToIn.

### Robustness of grouping

The grouping pattern by clustering *and* clading in our study agrees well with those of the protein trees (Misof et al. 2014; Thomas et al. 2020) except for Hemiptera group (see Notable differences in grouping and phylogenic positions in Discussions and Implications). Thus, it is surprising that the demographic grouping pattern is robust, in general, regardless of not only the information type (select protein-characteristics, or whole-proteome characteristics), but also of the methods (clustering or clading) used in grouping. For an implication of this result, see Similarities in grouping patterns in Discussion and Implications. However, not surprisingly, there are significant differences from the protein trees (Misof et al. 2014; Thomas et al. 2020) in branch-length and branching order of the groups (see Dissimilarities in branching orders and branch-lengths in Discussions and Implications).

### B. Emergence of the “Founders” of all major groups in a staged “burst”

For the following results we define “Cumulative Genomic Divergence (CGD)” for an internal node of the ToIn as the cumulative scaled-branch-length from the tree root to the node (see Cumulative Genomic Divergence (CGD) as “Evolutionary Progression Scale” in Materials and Methods) to represent the extent of the “evolutionary progression” of the node. The progression is scaled such that the root node of ToIn is set at CGD = 0 (see Outgroup in Discussions and Implications) and the leaf nodes of the extant organisms at CGD = 100, on average.

#### “Arthropodal burst” near the root of ToIn

Figures 3, 4 and 5 show the whole-proteome tree with CGD values. They reveal that the “founders” (for definition, see Supplemental information, Fig. S3) of all major groups of insects as well as non-insect arthropods (at Subphyla and Order levels) emerged in a staged burst within a short evolutionary progression span between CGD of 1.6 and 5.8 (marked by a small red arc in Figs. 3A and 3B), near the root of the tree. This observation is dramatically different from those of the protein trees (Misof et al. 2014; Thomas et al. 2020), where the founders of the major groups of all arthropods emerged throughout a long time-span of chronological scale.

**Fig 3A:**
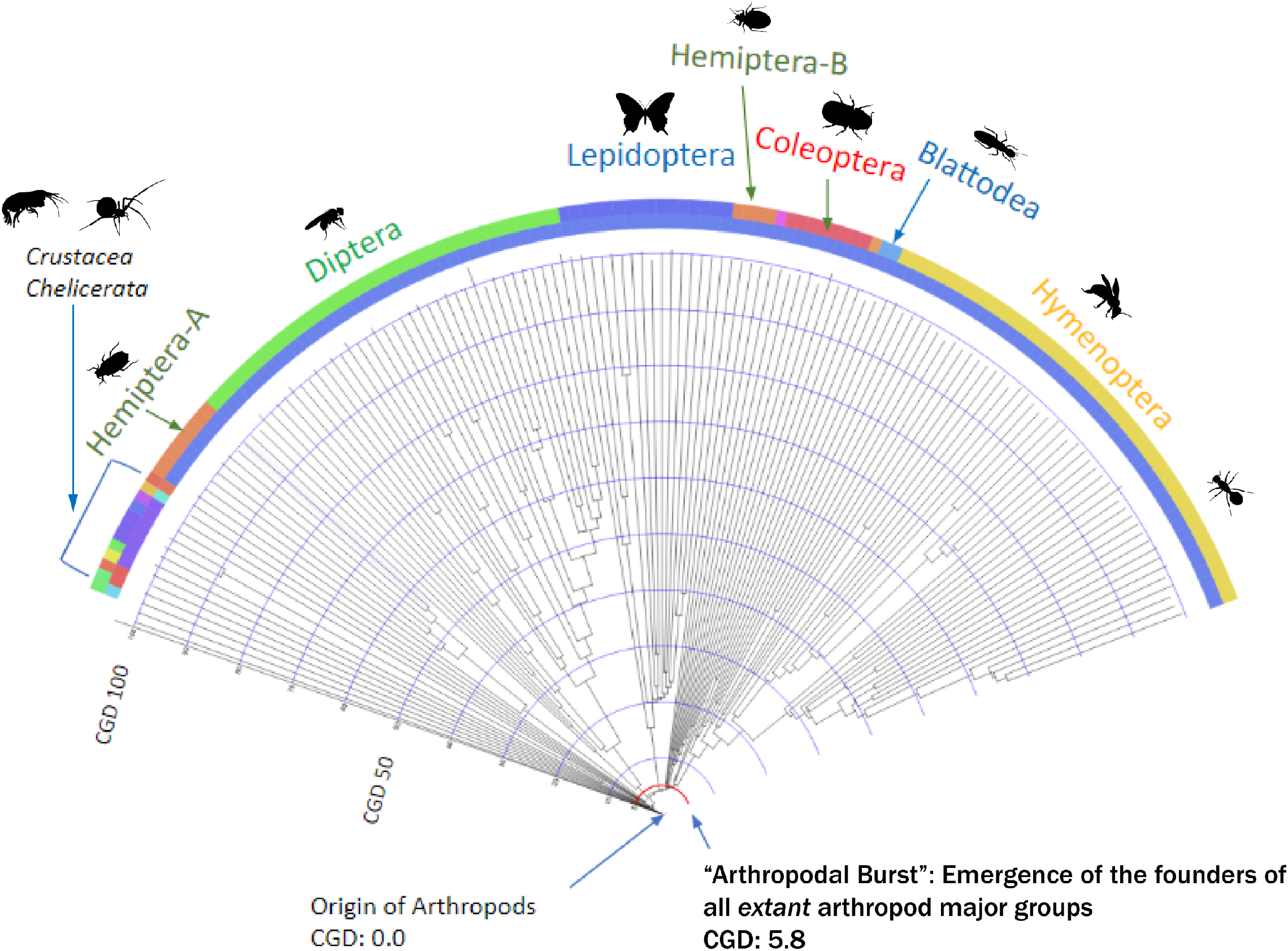
“Pie” representation of whole-proteome ToIn with the cumulative branch-lengths scale. This view of the whole-proteome ToIn shows all branch-lengths to emphasizes the progression of evolution of each member in the study population from the root of the tree at CGD = 0 to the extant forms of the members at CGD = 100, on average. The small red arc near the root is at CGD=5.8, by which point of the evolutionary progression the founders of all major groups (consisting of 7 Order groups and 2 Subphylum groups shown in Fig. 1) have emerged, suggesting that the remaining 94.2 on CGD scale corresponds to further diversification and gradual evolution of the founders and common ancestors *within* each major group toward their extant forms. The visualization of the ToIn was made using iTOL (Letunic & Bork 2019).

**Fig 3B:**
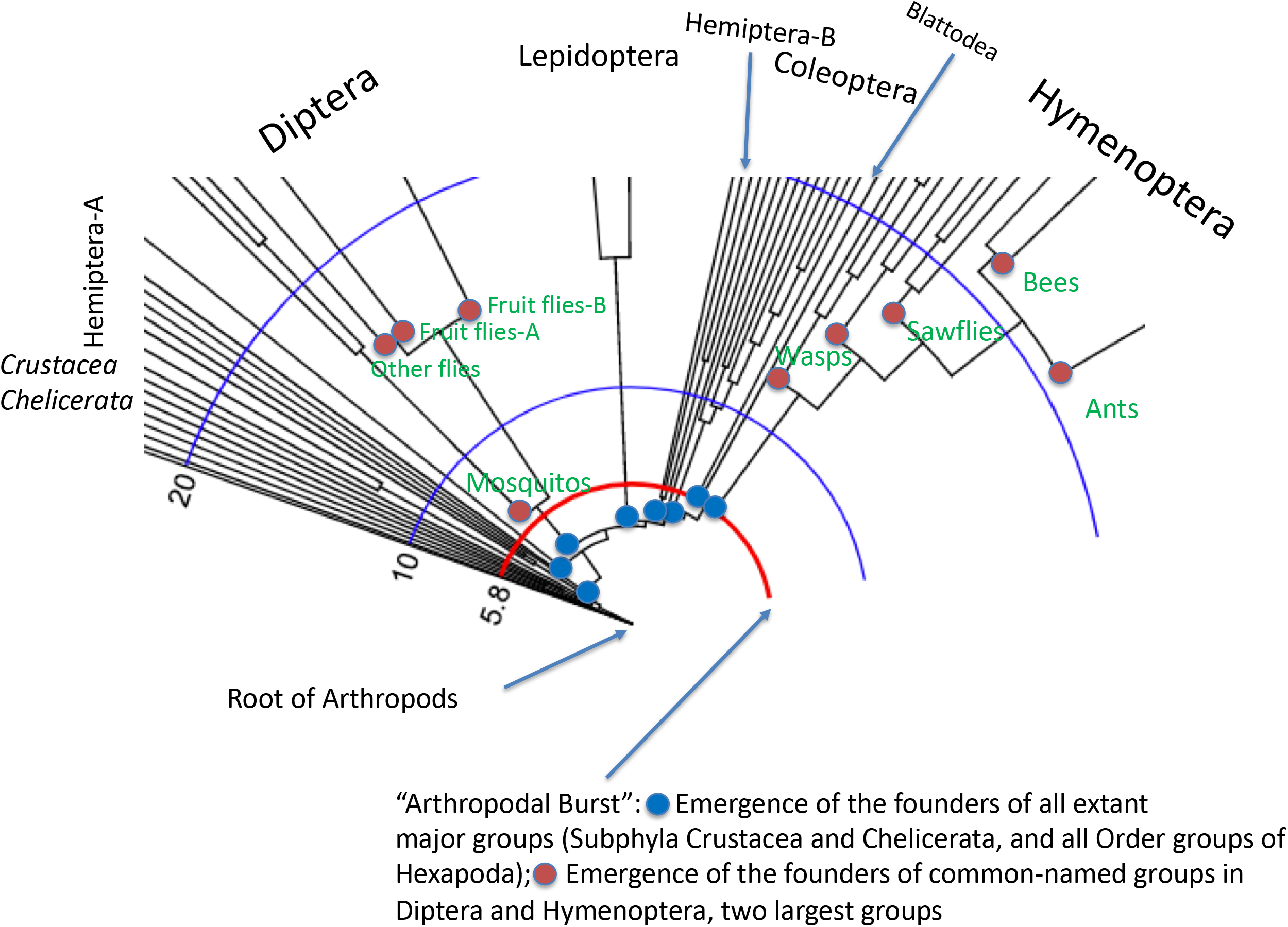
Expanded view of Fig. 3A near the root of the whole-proteome ToIn. Examples of the founders of all major groups are shown as blue dots, and the common ancestors of extant groups within two major groups, Diptera and Hymenoptera, as red dots. The visualization of the ToIn was made using iTOL (Letunic & Bork 2019).

**Fig 4:**
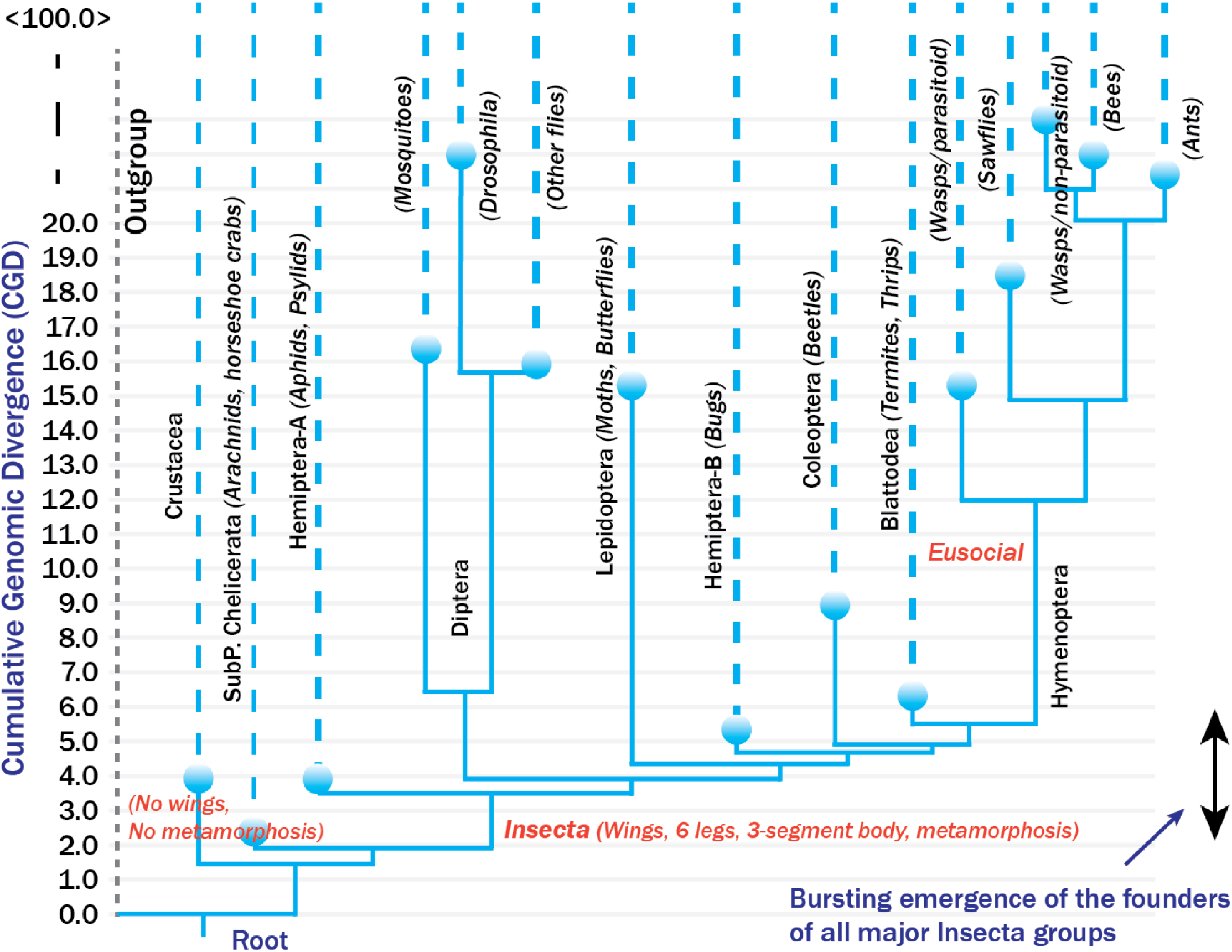
Simplified whole-proteome ToIn. The vertical axis shows cumulative genomic divergence (CGD) values, which ranges from zero to around 100, and they correspond to the extent of evolutionary progression from the root of the ToIn to the extant leaves. For simplicity, “singletons” (that do not belong to any named groups) are not shown, and all the leaf nodes and their branches of a common-named group (in parenthesis) are combined into a single dotted line coming out from their common ancestor node of the extant group shown as a blue sphere. Each internal node represents a “pool of founding ancestors” (see Supplementary Information Fig. S3). Dotted vertical lines are to indicate that they are arbitrarily shortened to accommodate large jumps of CGD values within a limited space of the figure. The double-headed arrow at bottom right indicates the short range of the CGD values, within which the founders of all the major groups of the extant organisms in this study have emerged in a “burst”. For our interpretation of horizontal lines and vertical lines, see Supplementary Information Fig. S3.

#### A subgroup of Hemiptera (Hemiptera-A) is the most basal group of all Insecta

The first founders of Class Insecta to emerge is the founders of Hemiptera-A group (aphids and a psyllid) at CGD of about 3.7 (Figs. 3A, 3B). This is in stark contrast to the protein trees, where all Hemiptera is the sister to Thysanoptera (thrips) (Misof et al. 2014) or a group of all Hemiptera, thrips and human louse is the sister to all other large groups of insects except Blattodea group (Thomas et al. 2020) (see also Notable differences in grouping and phylogenic positions in Discussions and Implications).

#### Order of emergence of the “founders” of all major groups of Insecta

Figure 4 shows a series of staged emergence of the founders of all major groups of Insecta. After the most basal group of Hemiptera-A group (aphids and a psyllid) at CGD of around 3.7, the founders of Diptera group emerged at CGD of 4.1, and those of the remaining five Order-level groups (Lepidoptera, Hemiptera-B (bugs, a planthopper and a whitefly), Coleoptera, Blattodea + a thrips, and Hymenoptera groups) at CGD of 4.4, 4.8, 5.2, 5.8, and 5.8, respectively. For possible implications see Notable differences in grouping and phylogenic positions in Discussions and Implications below)

## Discussions and Implications

### Similarities in grouping patterns

As mentioned earlier, it is surprising that the grouping patterns at Order level between the protein trees (Misof et al. 2014; Thomas et al. 2020) and our whole-proteome ToIn are very similar (see below for one notable exception of Hemiptera) despite the facts that the types of input data (multiple-aligned regions of selected proteins vs. whole-proteome) and estimation methods for evolutionary distance used (based on point mutational rates vs. whole genomic divergences) are very different. A possible implication is that, after the “burst”, the members of each group evolved largely “isolated” within the group without significant genomic mixing between the groups, thus, resulting in much smaller genomic variation within the group than between the groups, as manifested by mostly isolated clusters.

### Dissimilarities in branching orders and branch-lengths

It is not surprising that the branching orders and branch-lengths are not similar between the protein trees (Misof et al. 2014; Thomas et al. 2020) and our whole-proteome tree, because the assumptions under which the estimations for evolutionary distances among the organisms are calculated are very different: in the protein trees, the distances are calculated only for the aligned portions of the selected genes using, e.g., point-substitutional mutation rates, while in our tree they are calculated by accounting presence/absence of all amino acid short strings, Features, for all proteins due to all types of mutations.

### Evolutionary progression scale vs. Chronological time scale

It is difficult to design a scale that quantitatively measures the degree of evolutionary progression, because it is not clear what characteristics of an organism can best reflect the progression and also are quantitatively measurable. Since we are using whole-proteome sequence to represent each organism, we use the divergence of the whole-proteome sequences as the evolutionary progression scale (Choi & Kim 2020). In contrast to linear chronological time scale, the evolutionary progression scale is most likely not strictly linear, because any significant geological and ecological events may accelerate or decelerate the evolutionary progression for a given organism. However, the direction of arrows in both scales are the same, suggesting that the two scales may be calibrated when sufficient fossils, other independent records, and improved calibration methods become available (see Cumulative Genomic Divergence (CGD) as “Evolutionary Progression Scale” in Materials and Methods). Meanwhile, we use the evolutionary progression scale to compare the order of emergence of the founders of various major groups under the assumption that the whole-proteome sequence divergence can be considered as informational entropy, which increases as evolution progresses, similar to the physical entropy of universe increases as the universe evolve.

### “Burst” vs. Gradual emergence of the founders of major groups

While cognizant of the difference and similarity of the two scales, the most dramatic difference is observed in the span of the scales within which the founders of all major groups at Order level emerged in the protein trees (Misof et al. 2014; Thomas et al. 2020) and in our whole-proteome ToIn: In the protein trees, the founders of all the groups at Order level emerged gradually during a long chronological time span of about 350 Million years (Myrs) corresponding roughly 60% of about 570 Myrs between the tree root to the extant arthropods (Fig. 1 of Misof et al. 2014), or about 210 Myrs corresponding to roughly 37% of the same full chronological scale (Fig. 2 of Thomas et al. 2020). In drastic contrast, the founders of all the major groups in our tree emerged within about 4% of the full evolutionary progression scale in a sudden burst (“Arthropodal burst”; see Figs. 3A and 3B) near the root of our whole-proteome tree. This drastically contrasting observations between the two types of trees may have an important qualitative evolutionary implication in constructing the narrative for the birth of the insect diversity.

### Notable differences in grouping and phylogenic positions

Despite the drastic difference in the emergence pattern of the founders (burst vs. gradual) mentioned above, the order of emergence of the major groups at Order and Subphylum levels agree between the two protein trees (Misof et al. 2014; Thomas et al. 2020) and our whole-proteome tree with some notable differences in Hemiptera and Blattodea as described in Results above. These differences may get resolved once the whole-genome sequences of many more relevant organisms become available. At present, we suggest some possible implications as described below:

### Hemiptera

As mentioned earlier, in our whole-proteome ToIn (Figs. 3A and 3B) as well as in PCA (Fig. 1) and t-SNE (Supplemental Fig. S1) clustering plots, Hemiptera is divided into two separate clades/clusters, which we call Hemiptera-A (“primitive” Hemiptera, such as aphids and a psyllid) and Hemiptera-B (“bugs” such a planthopper, a whitefly, a stink bug and a bed bug), and their phylogenetic positions are very far apart (see Figs. 2, 3 and 5): Hemiptera-A at the basal position of all Insecta and Hemiptera-B as sister to the group consisting of Lepidoptera, Coleoptera and Hymenoptera. But, in the protein trees (Misof et al. 2014; Thomas et al. 2020) the both groups form a single clade, and is at basal or sister to all other large groups of insects except Blattodea group. This difference in clustering and phylogenetic positioning suggest that, when viewed at whole-proteome level, which includes *both* homologous and non-homologous proteins, the members are more similar within each subgroup than between the two subgroups in our tree. But, when viewed, as in the protein trees, only for the select homologous proteins in the *absence* of the non-homologous proteins, which are the overwhelming majority of the all proteins, they are similar among all of them to form only one clade.

### Blattodea

Two termites (Blattodea) and one thrips (Thysanoptera), both eusocial and hemimetabolous, form a clade in our whole-proteome ToIn and the clade is sister (or basal) to Hymenoptera group, which is also eusocial but holometabolous (see Fig. 5). However, in one protein tree (Misof et al. 2014), Blattodea group (cockroaches and termites, which are eusocial and hemimetabolous) is a member of a larger clade Polyneoptera and placed at the basal position to all other Order groups of Insecta, which are largely non-social and hemi- or holo-metabolous, while, in the other protein tree (Thomas et al. 2020), Blattodea group forms a separate clade, and is placed near the basal position of all other Order groups of Insecta. This is in contrast to what we observe in our ToIn, where Hemiptera-A, is the basal group of Insecta.

### Outgroup

Since our method does not require multiple sequence alignment, we constructed, as was described in our earlier works on whole-proteome trees (Jun et al. 2010; Choi & Kim 2017; Choi & Kim 2020), the proteome sequence of an “artificial (faux) arthropod” by “shuffling” (Knuth 1973; Fisher & Yates 1948) the alphabets of the whole proteome sequence of an organism in the study group. We used two such artificial arthropods (named R28612 and r12957) to form the outgroup for this study. Each has the same size and amino acid composition of corresponding protein of an extant arthropod, but does not have gene sequences information for the organism’s survival.

## Materials and Methods

### Sources and selection of proteome sequences

We downloaded the proteome sequences for 134 arthropods from NCBI RefSeq DB using NCBI FTP depository (O’Leary et al. 2016). Protein sequences derived from all organelles were excluded from this study. Also excluded from our study are those derived from whole genome sequences assembled with “low” completeness based on two criteria: (a) the genome assembly level indicated by NCBI as “contig” or lower (i.e. we selected those with the assembly levels of ‘scaffold’, ‘chromosome’ or ‘complete genome’), and (b) the proteome size smaller than 80% of the smallest proteome size among highly assembled arthropod genomes (*Anopheles gambiae str. PEST* with 14,089 proteins at “chromosome” assembly level; TaxID 180454).

All taxonomic names and their taxon identifiers (TaxIDs) of the organisms in this study are from NCBI taxonomy database, and listed in Supplementary Information, Dataset S1.

### Construction of whole-proteome Tree of Insects

Based on our earlier experiences of constructing whole-proteome trees of prokaryotes (Jun 2010), fungi (Choi & Kim 2017) and all life forms (Choi & Kim 2020) by Feature Frequency Profile (FFP) method (Sims et al. 2009), following choices have been made to obtain a topologically stable whole proteome ToIn of maximum parsimony (minimum evolution) by BIONJ (Saitou & Nei 1987): a) Among three types of genomic information (DNA sequence of the whole genome, RNA sequence of whole transcriptome and amino acid sequence of whole proteome) whole-proteome trees are most “topologically stable” as estimated by Robinson-Foulds metric (Robinson &Foulds 1981) at respective “optimal Feature-length”; b) For FFP as the “descriptor” of the whole proteome of each organism, the optimal Feature-length is about 10 amino-acid string (see Supplementary Information, Fig. S2); and c) Jensen-Shannon Divergence (JSD) (Lin 1991) is an appropriate measure of “divergence of information content”, as the measure of dissimilarity between two whole-proteome descriptors, for constructing the distance matrix of BIONJ (Saitou & Nei 1987; Gascuel 1997). It is important to note that such FFP of a whole-proteome sequence of an organism has all the information necessary to reconstruct the original whole proteome sequence.

### “Cumulative Genomic Divergence (CGD)” as “Evolutionary progression scale”

In Information Theory (Shannon 1948), the Jensen-Shannon Divergence (JSD) (Lin 1991), bound between zero and one, is commonly used as a measure of the dissimilarity between two probability distribution of informational features. The FFP as the descriptor for a linear sequence information of the whole proteome of an organism is such a probability distribution. Thus, a JSD value of two FFPs, used as a measure of the information divergence between two proteome sequences, is also bound between 0 and 1, corresponding to the JSD value between two FFPs of identical whole proteome sequences and two completely different whole proteome sequences, respectively. Any whole proteome-sequence “dissimilarity” between two extant organisms accumulated during the evolution can be considered as caused by changes of, ultimately, genomic sequences of all protein coding genes due to all types of mutational events, such as point substitutions, insertion/deletion of various lengths, inversion, recombination, loss/gain of genes, etc. as well as other unknown mechanisms, and they will bring JSD somewhere between 0.0 and 1.0 depending on the degree of the sequence divergence.

In this study the collection of the JSDs for all pairs of the study organisms plus 2 out-group members (see Outgroup in Discussions and Implications) constitutes the “distance matrix” for BIONJ (Saitou & Nei1987; Gascuel 1997). Since all the branch-lengths are derived from the JSD values, the cumulative branch-length of an internal node, which we call “cumulative genomic divergence (CGD)” (to reflect the fact that the proteomic divergence is ultimately derived by the genomic divergence during evolution) of the node, can be considered as the point of evolutionary stage reached by the node on an “evolutionary progression scale”. For convenience of assigning the nodes on the progression scale, CGDs are scaled, as mentioned earlier, such that the CGD value at the root node of ToIn is set to zero and the leaf nodes of the extant organisms to 100, on average, corresponding to the fully evolved genomic states of the organisms, which we define as the beginning and ending point of the “evolutionary progression scale” for the organisms (see Fig. 3A).

### Clustering methods

We use two unsupervised methods to observe the clustering patterns based solely on whole-proteome sequences: Principal Component Analysis (PCA) and t-Distributed Stochastic Neighbor Embedding (t-SNE) (R Core Team 2016; v.d. Maaten & Hinton 2008). Both are dimensional reduction methods, but with different strengths and weaknesses for our purposes, which help to visualize any clustering pattern in the data distribution. Both are based only on the evolutionary distances (CGD in this study), estimated by the divergence of whole-proteome sequences among all pairs of the study arthropods. In PCA, the distances within a cluster as well as between two clusters are quantitative, thus, two close clusters nearby may not resolve well. In t-SNE, which applies Machine Learning to emphasize the resolution of nearby clusters, but the inter-cluster distances are de-emphasized, thus, not quantitative.

## Supporting information

Supplementary Information.ISE.V2.pdf

## Declarations

## Acknowledgement

We gratefully acknowledge the comments and advices from the arthropod experts at University of California, Berkeley, CA., Profs. Kipling Will and Peter Oboyski, and the constructive suggestions on our application of Information Theory and Unsupervised Clustering methods from Prof. Se-Ran Jun of College of Medicine, University of Arkansas for Medical Sciences, Little Rock, AR, and Dr. Chao Zhang, Chief Scientific Officer, of Plexxikon, Inc. All the silhouettes are generated by Rigel Sison.

## Author contribution

Conceptual design of the study and speculative interpretations and implications of the results by SHK; filtering and curation of genomic and proteomic sequence data from NCBI database, computational-algorithm design, programming and execution by JJC and BJK; unsupervised clustering by various algorithms were performed by BJK; interpretation of computational results by SHK, JJC and BJK; manuscript preparation by SHK with extensive discussions with JJC and BJK; all figures are designed by SHK, JJC and BJK.

## Computer code availability

The FFP programs for this study (2v.3.0) written in GCC(g++) is available in Github: https://github.com/jaejinchoi/FFP.

