## Supplementary Information.ISE.V2.pdf for "Whole-proteome Tree of Insects: Grouping and phylogeny without sequence alignment"

**Supplementary data:** Dataset S1.xlsx

### Legends for Supplementary Information Figures

**Figure S1. Two-dimensional plot of t-Distributed Stochastic Neighbor Embedding (t-SNE)**
t-SNE (v.d. Maarten & Hinton 2008) is a machine learning algorithm of clustering by reducing a high-dimensional data into a two or three dimensional space for easy visualization by emphasizing resolution of clusters, but de-emphasizing the distances between the clusters. There are about 17 clusters with their common names in parentheses. Most of the clusters in classical PCA (Fig. 1) are also observed in t-SNE plot including the split Hemiptera into two separate clusters. These clusters can be assorted into six Order groups and two Subphylum groups in colored bold-letters, correspond to Hemiptera (dark green), Lepidoptera (light green), Diptera (blue), Coleoptera (pink), Blattodea (yellow), Hymenoptera (light blue), Chelicerata (gray and red) and Crustacea (gray). Some clusters are loose, and there are a few unclustered organisms in gray.

**Figure S2. Selection of the tree with most stable topology as a function of Feature length.**
Topological variation, as represented by Robinson-Foulds metric (Robinson & Foulds 1981) between a pair of trees, is shown on Y-axis, and Feature length on X-axis. Robinson-Foulds metric is calculated between two trees: one for Feature length of $l$ and the other for Feature length of $l+1$. This figure shows three types of genomic information (DNA sequence of whole genome, RNA sequence of whole transcriptome and amino acid sequence of whole proteome). Of the three, the whole proteome sequence-based genome ToIn is most topologically stable, reaching the lowest point of the plot, starting from the Feature length of around 8 and remain stable for longer Feature lengths. For this study we use Feature length of 10.

**Figure S3. Internal node as a Pool of founding ancestors**

As described in our earlier work (Choi & Kim 2020), an internal node in this study (shown as a rectangle) is represented as “a pool of diverse founding ancestors” like a bucket of “mosaic tiles”, not as a “clonal single tile”. Each internal branch is divided into two components: the horizontal arrow represents the *emergence* of *one or a small subset of founding ancestors* in the pool under “abrupt” evolutionary event with drastic environmental and ecological changes, and the vertical arrow, which represents the genomic *diversification* of the emerging founder to evolve into a new pool of founding ancestors through relatively *gradual* evolution or a “common ancestor” of a clade, shown as a circle, which is the last internal node containing all extant members of a named clade of arthropods.

Suppl. Fig. S1

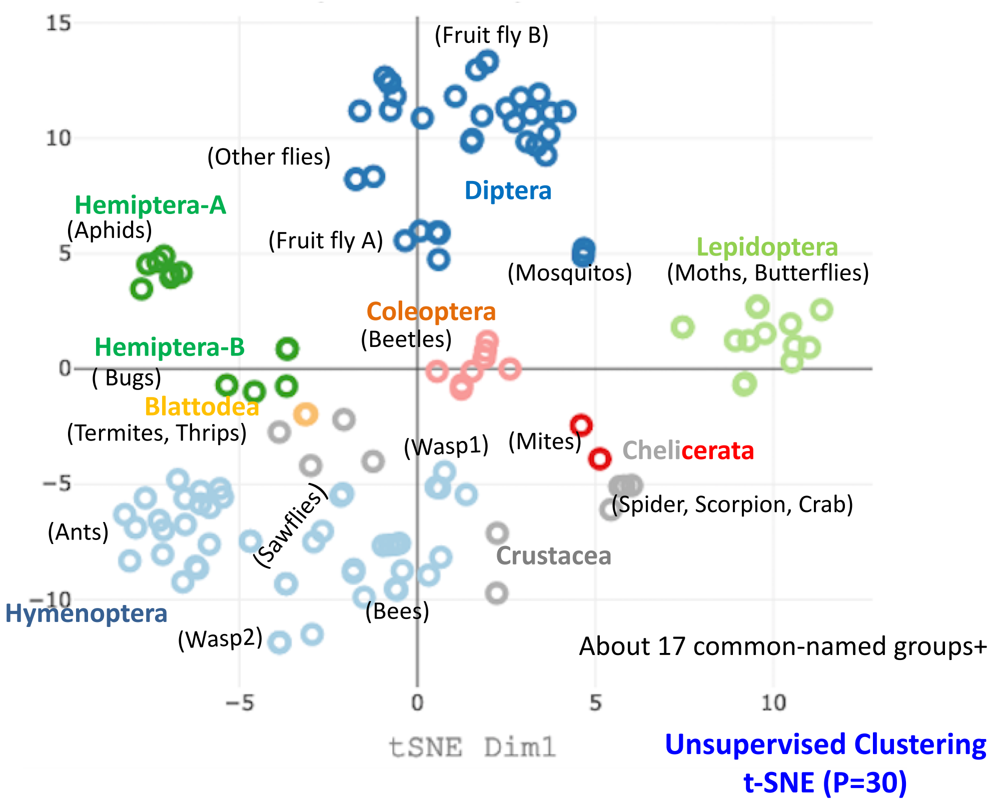

Suppl. Fig. S2

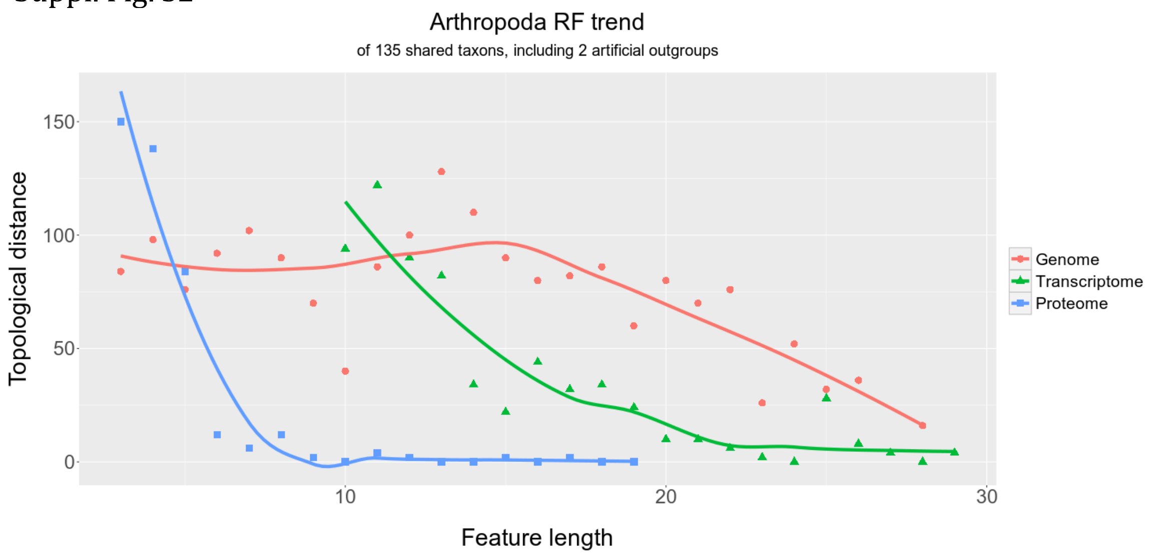


Suppl. Fig. S3

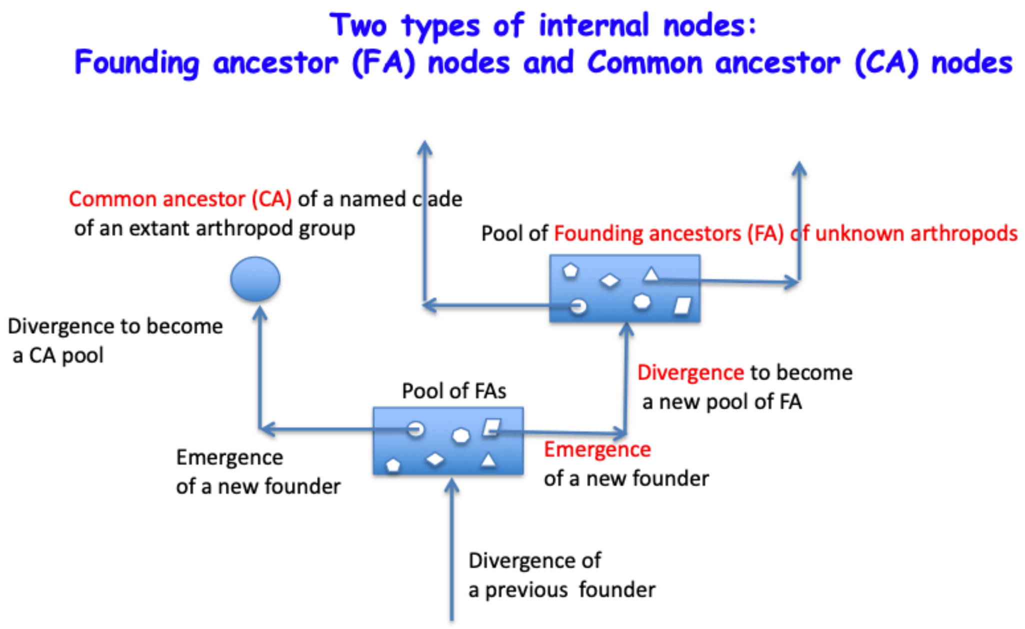
